# The pod components of the *Shigella* T3SS sorting platform accommodate multiple copies of Spa33 (SctQ)

**DOI:** 10.64898/2026.02.26.708312

**Authors:** Sean K Whittier, Shoichi Tachiyama, Samira Heydari, Wendy L. Picking, Jun Liu, William D. Picking

**Affiliations:** Department of Veterinary Pathobiology, University of Missouri, Columbia, MO 65211; Department Christopher S. Bond Life Sciences Center, University of Missouri, Columbia, MO 65211; Department of Microbial Pathogenesis, Yale University West Haven, CT 06516; Microbial Sciences Institute, Yale University, West Haven, CT 06516

**Keywords:** *Shigella*, type III secretion system, injectisome, Spa33, SctQ, sorting platform

## Abstract

The bacterial type III secretion system (T3SS) uses a membrane-embedded injectisome assembly to export effector proteins into host cells. While atomic-level structural details have been revealed for much of the T3SS apparatus, the model of the cytoplasmic sorting platform remains largely low-resolution. A central structural element of the sorting platform is the so-called "pod" protein, SctQ, which anchors the sorting platform to the inner membrane via interaction with the adaptor protein SctK, and connects to the central ATPase via the spoke protein SctL. SctQ proteins also interact with alternatively translated homodimers of their C-terminal SPOA2 domains. Low resolution electron density maps have provided an outline of the sorting platform architecture, and fluorescence microscopy studies have suggested a 1:4:2 SctK:SctQ:SctL stoichiometry. While there are experimental and AlphaFold structures of the individual components or complexes of sorting platform pod, there is currently no model for the pod structure that adequately fits the electron density or accounts for the proposed stoichiometry. Here we use AlphaFold to generate a model of the *Shigella* pod complex in which two copies of the SctQ protein, Spa33, bind the adaptor protein MxiK, with each copy of Spa33 bound to an alternately translated SPOA2-SPOA2 domain. We show through mutation of energetically critical interface residues, predicted by computational mutant scanning, that both Spa33 binding sites on MxiK are required for T3SS activity in *Shigella flexneri*, as well as binding of Spa33 to the SPOA2-SPOA2 homodimer. We find that this model fits well to the upper two-thirds of the pod electron density, albeit in a manner that places the protein components slightly closer to the inner membrane than traditionally presented. Further, cryogenic electron tomography of mutant injectisomes reveals the lack of a complete sorting platform and may suggest an alternative model featuring four copies of Spa33 that still fits the upper pod electron density but better explains unfilled density in the lower third of the pod.

## Introduction

The type III secretion system (T3SS) is used by gram-negative bacterial pathogens to secrete protein effectors into eukaryotic host cells^1^. These effectors are secreted by the type III secretion apparatus (T3SA), or injectisome, the architecture of which is conserved among phylogenetically distant pathogens^2^. The T3SA can be divided into three distinct portions (see Suppl. Fig. 1): a hollow needle-tip assembly that extends from the outer membrane to contact host cells, basal body embedded in the inner and outer membranes, and a cytoplasmic sorting platform that is responsible for substrate recognition and driving secretion^3–5^. Given its role in human disease, elucidating the structure of the T3SA is critical from the standpoint of developing drugs that inhibit its function or vaccines that may prevent severe disease. The T3SA is also a complex protein nanomachine, and the study of its structure and function may allow us to glean basic biophysical and biochemical principles that govern large biomolecular assemblies.

Perhaps the least understood portion of the T3SA is the cytoplasmic sorting platform^6^, this highly dynamic macromolecular complex is thought to be transiently assembled, with some individual protein components in frequent exchange with the cytosol^7,8^. Its overall organization is understood: a central ATPase (SctN in unified nomenclature^9^) is connected via radially symmetric spokes (SctL) to six "pods”, formed by SctQ, that then connect to membrane-bound ring of SctD via the adaptor protein, SctK^5,10^. The SctQ protein is comprised of independently folding N- and C-terminal domains^11^, SctQ_N_ and SctQ_C_. The structures of the C-terminal domains of SctQ proteins from *Shigella* (Spa33) and *Salmonella* (SpaO) have been shown experimentally to be a dimer of two "surface presentation of antigens" (SPOA) domains, SPOA1 and SPOA2, and NMR data show that SpaO_C_ binds to a peptide from the SctL spoke protein OrgB^11,12,13^. Furthermore, SctQ genes feature an internal translation start site upstream of SPOA2^14^. These alternatively translated SPOA2 domains form a SPOA2-SPOA2 homodimer, SctQ_short_, which associates with the full-length SctQ protein and may be required for T3SS function.

Recent insight into the pod structure has come from AlphaFold modeling and *in vivo* crosslinking of the SctQ complexes with SctK, SctL, and SctQ_short_ in the *Salmonella* system^15,16^. Those results show that SctQ interacts with SctK via its N-terminal domain, and with SctL through SctQ_C_. Additional work reported the interaction between SctQ and SctQ_short_ and showed that SctK simultaneously binds to SctL, SctQ, and SctQ_short_. While those results have greatly expanded our understanding of the organization of the pod, a single SctK:SctQ:SctL complex is not sufficient to fill the electron density map from cryo-electron tomography (cryo-ET)^15^. Furthermore, fluorescence microscopy of the *Yersinia* injectisome indicates that each pod should contain as many as four copies of SctQ and two copies of SctL^7^. A full model of the sorting platform, therefore, should include multiple copies of SctQ and SctL, fit well to the electron density map, and retain the molecular interactions revealed in experimentally verified AlphaFold models.

In this work, we present a model for the upper two-thirds of the *Shigella* sorting platform pod. We show that AlphaFold predicts a complex of the *Shigella* SctK protein MxiK binding two copies of the SctQ protein Spa33 with high confidence and demonstrate, through mutation of critical interface residues, that each of these binding sites on MxiK is required for T3SS function in *Shigella flexneri*. Additionally, we find that mutations to the AlphaFold-predicted Spa33/Spa33_short_ interface drastically reduce or eliminate T3SS activity, indicating that Spa33_short_ is a critical component of the pod.

The AlphaFold-generated complex of MxiK with two copies of full-length Spa33 and two copies of Spa33_short_ fits well into the upper portion of pod density in cryo-ET maps of the *S. flexerni* sorting platform, although the protein components are shifted “upward” toward the inner membrane relative to where they previously have been shown to reside in the cryo-ET density map (Suppl. Fig. 1). This positioning is somewhat non-canonical, but it fits the density well and explains the curious hole seen between regions commonly attributed to MxiK and Spa33 (Fig. 3c).

Finally, cryo-ET structures of pod complex mutants retain some pod density but lack density for the SctL spoke MxiN and central ATPase, Spa47. The presence of pod density in these mutant injectisomes suggests that a stable pod structure is required for recruitment of the ATPase. It may also suggest a possible alternative model comprising four copies of Spa33 that may explain additional density at the bottom third of the pod not accounted for by the MxiN dimer, as will be presented in the Discussion.

## Results

### Spa33 binds MxiK at two sites

AlphaFold predicts a similar structure as reported in the *Salmonella* T3SS when asked to predict the multimer structure of Spa33 and MxiK in a 1:1 stoichiometry (Suppl. Fig. 2a)^15^. The resulting model shows Spa33_N_ binding to a hydrophobic patch on the surface of MxiK. However, there is another, similarly sized hydrophobic patch near this binding site on the surface of MxiK (Suppl. Fig. 2b) that is not predicted to be occupied by either copy of the MxiG cytoplasmic domain. Further, a 1:1 Spa33:MxiK interaction model does not fully account for observed electron density in this region of the sorting platform. Therefore, we used AlphaFold to predict the structure of a 2:1 Spa33:MxiK complex. Here, the predicted model shows that one copy of Spa33_N_ binds to the same site as in the 1:1 complex but now has the second copy of Spa33_N_ binding to the adjacent, unoccupied hydrophobic patch on MxiK (Fig. 1a). AlphaFold accurately predicts the structure of each component protein domain, as indicated by pLDDT scores (Suppl. Fig. 2c), and is highly confident of the overall complex structure, with low PAE scores across all residues (Fig1b). pTM and ipTM scores of the predicted complex are 0.82 and 0.79, respectively, indicating that the prediction is likely that of the “true” structure. This 2:1 SctQ:SctK interaction is not unique to *Shigella*, as AlphaFold predicts similar 2:1 complex for SpaO/OrgA in *Salmonella* and YscQ/YscK and PscQ/PscK in *Yersinia* and *Pseudomonas*, with low PAE scores in each case (Suppl. Fig. 3).

**Figure 1.**
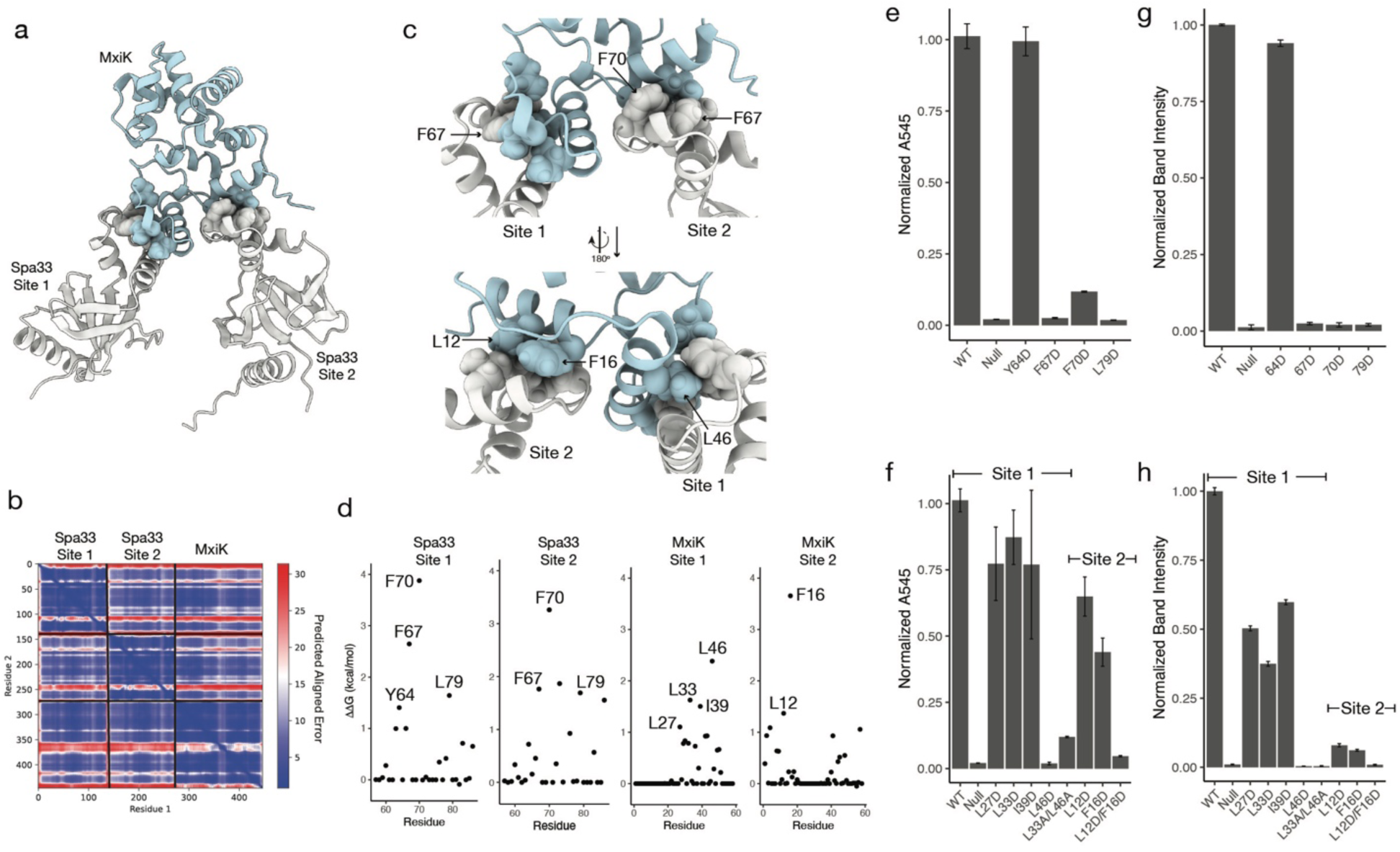
a) AlphaFold mutltimer structure of MxiK (blue) bound to two copies of *Spa33_N_* (white). *Spa33_C_* is omitted for clarity. Spa33 binding site 1 is homologous to the interaction reported between SpaO and OrgA in *Salmonella*. b) Plot of predicted aligned error for the Spa33_N_:Spa33_N_:MxiK complex. c) Close-up of Spa33/MxiK interaction sites, with energetically critical residues indicated by spheres. d) Results of computational alanine scanning of the Spa33_N_ and MxiK binding sites. Results of contact-mediated hemolysis for Spa33 and MxiK binding site mutants are shown in e) and f), respectively. The corresponding CR-induced secretion of IpaB is shown in g) and f), respectively. The quantification of IpaB band intensity is from western blots. All values are normalized to average of WT band intensity.

The predicted complex indicates that Spa33 binds to each site of MxiK using the same surface, specifically the region between residues 64-86, which form helices 3 and 4 of Spa33_N_ and the turn between them (Fig. 1 a and c). Computational alanine scanning of each Spa33 interface with MxiK reveals large energetic binding contributions from the same group of hydrophobic residues in Spa33_N_, with F67 and F70 predicted to be hot-spot residues (ΔΔG > 2.0 kcal/mol) (Fig 1d), and Y64 and L79 near the hot-spot cutoff^17^. To confirm that this surface is involved in MxiK binding, we made perturbing mutations at each of these positions, expressed the mutant proteins in a Δ*spa33* (Spa33_Null_) strain of *S. flexneri,* and characterized secretion phenotype using contact mediated hemolysis^18^ and Congo red (CR)- induced secretion^19^.

The replacement of each residue with aspartic acid, which places a formal charge in the hydrophobic interface, largely eliminates hemolytic activity for F67D, F70D, and L79D, with only F70D retaining a small level of hemolysis above that of the uncomplemented Spa33_Null_ strain (Fig. 1e). The Y64D mutant, however, retained wild-type hemolysis capability. These results are mirrored in the case of CR-induced secretion. F67D, F70D, and L79D all had no detectable IpaB or IpaD secretion, while Y64D secreted wild-type quantities of each (Fig. 1g, Suppl. Fig. 4). Single alanine substitutions did not greatly reduce hemolytic activity for any of these positions, however the simultaneous replacement of F67 and F70 with alanine resulted in a null phenotype (Suppl. Fig. 2d). Western blots were used to confirm that none of these mutants affected expression levels of either full-length Spa33 or Spa33_short_ (Suppl. Fig.5).

The two binding sites on MxiK, as mentioned above, are largely hydrophobic. Computational alanine scanning of MxiK interface residues reveals a handful of hot-spot residues that dominate energetically. At the first binding site, which is homologous to the interaction described in *Salmonella*, one hot-spot residue is predicted, L46, with three other residues, L27, L33, and I39, having ΔΔG > 1 kcal/mol upon mutation to alanine (Fig 1d). The second binding site is energetically dominated by F16, with F16A giving a predicted ΔΔG > 3.5 kcal/mol. Three other residues have ΔΔG > 1 kcal/mol, with L12 have the largest predicted contribution of these. As with Spa33, we confirmed these interfaces by mutating energetically critical residues, expressing the mutants in a Δ*mxiK* (MxiK_Null_) *S. flexneri strain*, and determining contact-mediate hemolysis and CR-induced secretion.

Single aspartate substitutions of binding site 1 residues L27, L33, and I39 had a minor impact on translocon pore formation, with each mutant having about 75% wild-type hemolysis activity. F46D, on the other hand, had no detectable hemolysis (Fig. 1f). Additionally, a double alanine mutant, L33A/F46A, was only 10% as active as wild-type MxiK (Fig. 1f). L27D, L33D, and I39D had larger impact on induced secretion, with each secreting 50% or less than wild-type. F46D and L33A/F46A each had no detectable IpaB or IpaD secretion (Fig. 1h, Suppl. Fig. 4).

Two residues at binding site 2 were mutated next (L12 and F16). Aspartate substitutions L12D and F16D were modestly perturbing, causing 40% and 65% loss of hemolytic capability relative to wild-type, respectively (Fig. 1f), while a simultaneous substitution of each residue with aspartate resulted in a null hemolysis phenotype (Fig. 1f). Again, mutant effects on induced secretion were greater than those seen in hemolysis. Both L12D and F16D showed ∼95% reduction in secretion compared to wild-type (Fig. 1h, Suppl. Fig. 4). The double mutant L12D/F16D had no visible IpaB or IpaD bands after CR-induction (Fig. 1h, Suppl. Fig. 4). As with Spa33, western blots show that these mutations do not impact MxiK expression levels (Suppl. Fig. 5).

### Spa33 binding of the Spa33_short_ homodimer is required for T3SS function

The *spa33* gene features an internal translation start site just upstream of the *SPOA2* region^11,14,20^. This results in an alternately translated SPOA2 protein product, which forms a SPOA2-SPOA2 homodimer, Spa33_short_^14^. Recent work using AlphaFold with the *Salmonella* Spa33 homologue, SpaO, has shown that the SPOA2 homodimer binds to the full length SpaO protein between the N- and C-terminal domains^16^. In the case of Spa33, AlphaFold predicts a similar complex with the Spa33_short_ homodimer binding along the flexible linker between Spa33_N_ and Spa33_C_ (Fig. 2a, Suppl. Fig. 6). The confidence of the AlphaFold structure is high, with low PAE values between domains (Suppl. Fig. 6).

**Figure 2.**
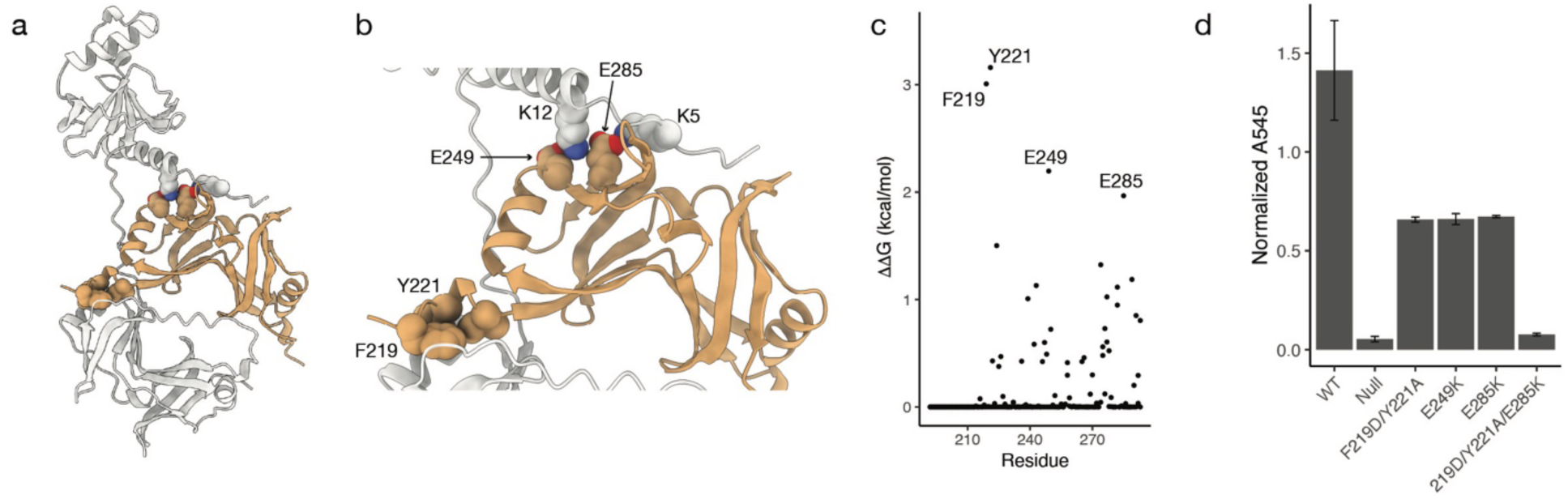
a) AlphaFold model of the Spa33/Spa33_short_ complex. b) A close up of interacting regions between Spa33 and Spa33_short_, with residues predicted to be energetically critical to complex formation indicated, is shown by red and purple spheres. c) Results of computational alanine scanning of Spa33_short_ homodimer residues are presented by the alanine scanning. d) Contact-mediated hemolysis shows a loss of hemolytic activity when critical Spa33_short_ residues are mutated.

Computational alanine scanning of the Spa33/Spa33_short_ complex reveals two predicted hot-spot residues, F219 and Y221, and several other residues predicted to have ΔΔG > 1 kcal/mol (Fig. 2c). The locations of these residues indicate two critical areas of interaction between Spa33 and the SPOA2 homodimer. The first of these involves F219 and Y221, along with I224, which insert into a hydrophobic pocket in Spa33_C_ (Fig. 2b). The second area of interaction is defined by two salt-bridges formed between E249 and E285 in Spa33_short_ and K12 and K5, respectively, in the N-terminal peptide of Spa33_N_ (Fig. 2b).

Again, we experimentally confirm the importance of these interactions through mutation and phenotypic characterization of T3SS activity using contact-mediated hemolysis. Because of their sequential proximity, F219 and Y221 were mutated together.

Mutation of the F219/Y221 positions or either of the salt-bridge residues in Spa33_short_ resulted in a greater than 50% loss of hemolytic function, relative to wild-type. Simultaneous mutation of F219, Y221, and E285, however, eliminates T3S activity (Fig. 2d). Western blots of *S. flexneri* overexpressing these mutants show levels of both full-length Spa33 and Spa33_short_ comparable those of wild-type Spa33 (Suppl. Fig. 5), indicating that these mutations do not affect protein expression. These results indicate that binding of Spa33 with Spa33_short_ is required for a fully functional T3SS in *Shigella*.

### Fitting model of upper pod region to experimental electron density map

The above results lead us to propose a model for the upper portion of the SP pod in which a single copy of MxiK is bound to two copies of Spa33, each of which is complexed with a Spa33_short_ homodimer (Fig 3a, b). If this model is valid, it should fit within experimentally obtained electron density maps of the *Shigella* SP. EM density fitting was performed by using ChimeraX^21^ to simulate EM density of our model at a resolution of 13 angstroms and then fit the simulated density to a density map from a focused refinement of the sorting platform structure by maximizing the cross correlation between them. As a sanity check for model fitting, we used AlphaFold to include two copies of the MxiG cytoplasmic domain (MxiG_C_) bound to MxiK, as well as a short peptide from MxiN bound to each copy of Spa33_C_. The fit model should position MxiG_C_ close to the inner membrane and the MxiN peptide near the spoke density.

The best fit of this model to the experimental electron density map is shown in Figure 3, panels d-f. This fit gives a correlation between simulated and experimental electron density maps of 0.74, and places MxiG_C_ near the inner membrane and the MxiN peptide in the density region close to the spoke. Importantly, the position of each protein component is shifted upward toward the membrane relative to the current placements in most models of the SP. The region of density commonly designated as arising from MxiK is occupied by two copies of Spa33_N_ in this model, with MxiK and the MxiG_C_ domains occupying a common region of density closest to the inner membrane. Attempts to fit our model into alternative positions in the density map give poor correlation values and do not visually appear to fit the density well.

The model presented in Figure 3d leaves the lower third of the pod density unoccupied. This density must arise, at least in part, by a dimer of the SctL spoke protein MxiN, which must be able to connect the central ATPase to Spa33_C_. Unfortunately, AlphaFold does not predict the structure of the MxiN dimer with high confidence and will often predict different structures for different seeds or when the Spa33_C_ binding partner is present (Suppl. Fig. 7). We are, therefore, unable to model this portion of the density. Assuming, however, that there is a bend in the MxiN dimer, such as that shown in the AlphaFold structure in Suppl. Fig. 7c, it is possible to connect the Spa47 ATPase to Spa33_C_. Fig. 3f shows a hypothetical conformation of the MxiN dimer that connects Spa33_C_ to Spa47. Here, each MxiN monomer extends downward from Spa33_C_ until it curves toward Spa47 around residue 57 and forms an extended coiled-coil alpha helix. This is not to propose that this is the actual conformation of MxiN, but only to show that the position of Spa33_C_ in Figure 3d is not too distal to attach to the spoke.

### Cryo-EM structures of pod mutant injectisomes

To understand how interfacial mutants affect the T3SA, we used cryo-ET to compare in-situ structures of mutant injectisomes to a non-secreting T3SA in which the Spa47 ATPase is inactive (Spa47_K165A_)^22^. Minicells of *Δspa33* or *ΔmxiK S. flexneri* complemented with *spa33*_F67A/F70A_, *spa33*_F70D_, *spa33*_F219D/Y221A_, or *mxiK*_L46D_ were prepared and visualized by cryo-ET. Sorting platforms were then compared through their averaged structures.

Prior to averaging, individual injectisomes in tomograms were classified according to the presence or absence of either extracellular needles or any indication of sorting platform density. Fewer than 1% of injectisomes had extracellular needles, indicating they are largely non-secreting. Classification of subtomograms based on sorting platform density is more difficult than the presence of needles. Here, we classified injectisomes with any visual indication of sorting platform density, no matter how faint, as sorting platform positive (SP+), and those without as sorting platform negative (SP-). Pod mutant injectisomes were less likely to be SP+ than the Spa47_K165A_ mutant (88% SP+), ranging from 65% SP+ (Spa33_F70D_) to 37% (Spa33_F219D/Y221A_).

Subtomogram-averaged structures also indicate that pod mutant injectisomes are inactive. The basal body of all mutant structures, and Spa47_K165A_, appear in a “closed” conformation, with no visible inner rod assembly and the presence of a septum structure blocking the channel (Fig. 4). Unlike Spa47_K165A_, which has a fully intact sorting platform, none of the averaged pod mutant structures have any density for Spa47 or the MxiN spokes. Interestingly, some pod density is apparent for each mutant sorting platform. Going by the model presented here in Figure 3, all Spa33/MxiK interaction mutants have regions of upper pod density corresponding to MxiK and Spa33, as well as density for the lower third of the pod. The density of the Spa33_short_ interaction mutant Spa33_F219D/Y221A_ is less resolved in the lower third of the pod. These data suggest that mutation of the Spa33/MxiK interface or of the Spa33/Spa33_short_ interface destabilizes the pod structure such that, while pods may still transiently form, they are unable to form spokes or recruit the Spa47 ATPase. Because of a sixfold radial symmetry operation performed on the data during processing, we cannot state precisely how many pods are present in the mutant sorting platforms, but the weaker pod density relative to Spa47_K165A_ suggests fewer than the full complement of six pods.

**Figure 3.**
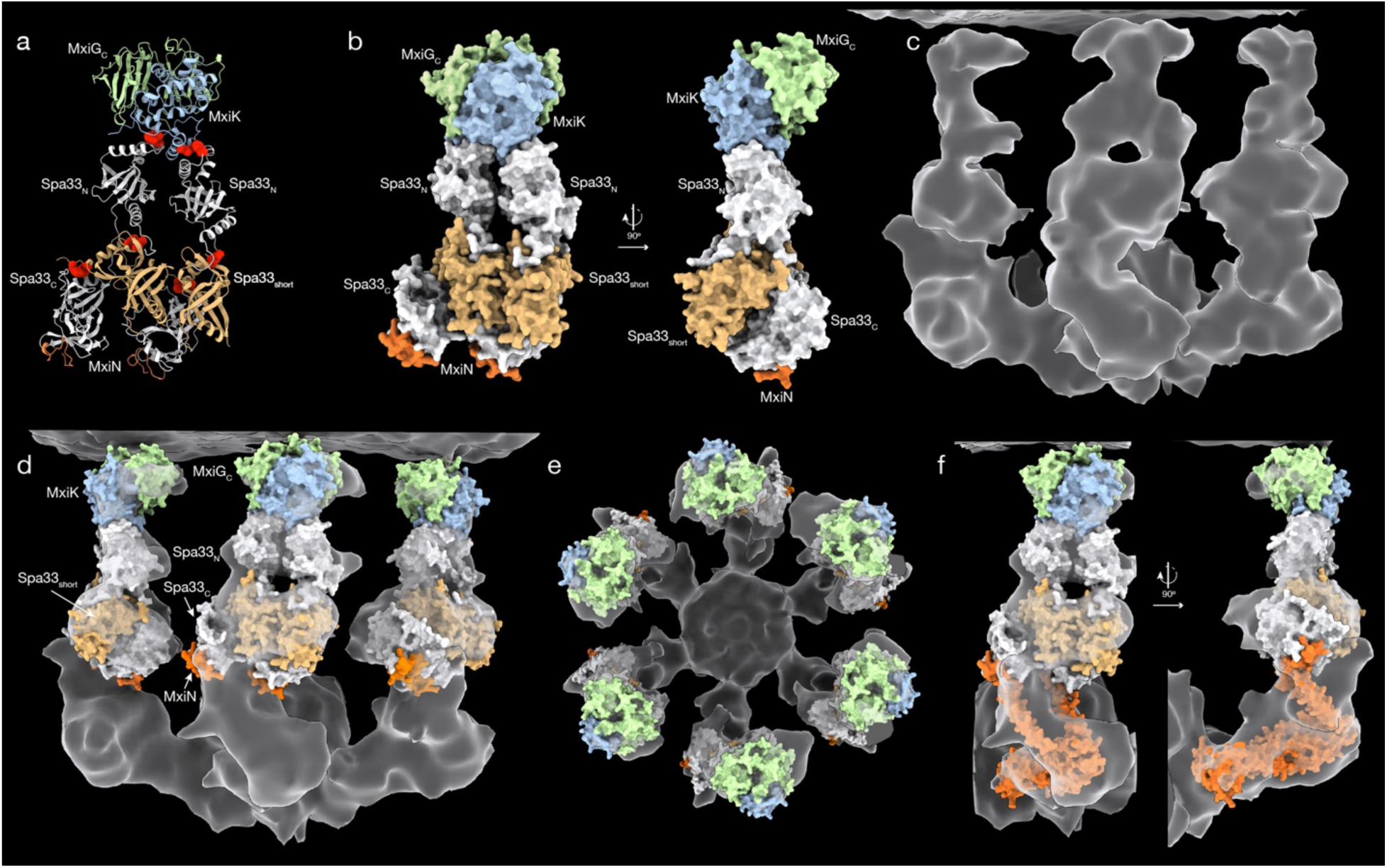
a) Cartoon representation of full model of MxiK/Spa33 pod complex. Positions of interface disrupting mutants are indicated by red spheres. The MxiG (SctD) cytoplasmic domains (green) and MxiN peptides (red) are included as a check on feasibility of model fitting to the electron density map of the sorting platform. b) Surface representation of the model shown in panel (a), including 90° rotated view. c)The experimental electron density map of *S. flexneri* sorting platform is shown. d) A fit of the pod complex model to the electron density map of the *Shigella* sorting platform. The inner membrane is at the top of the map and the MxiN spokes and central ATPase are at the bottom (intervening MxiG cytoplasmic domains are not shown). Likewise, the back three pods have been cropped out for clarity. e) A view of the model fit is shown looking down from the inner membrane. f) A hypothetical model of a MxiN dimer (red) is fit into electron density to show that the spoke can still connect the ATPase with Spa33_C_ (white) in this new model.

**Figure 4.**
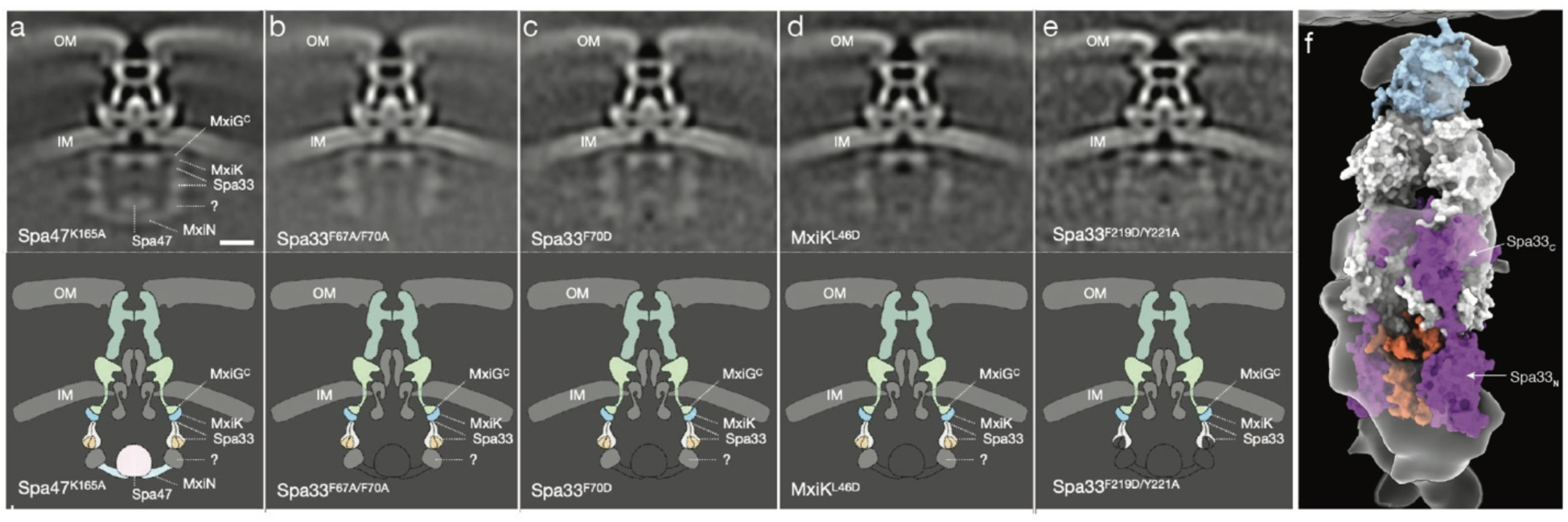
a-e) Subtomogram averaged structures of injectisomes from *Shigella* minicells expressing an inactive variant of the ATPase Spa47 (Spa47_K165A_, positive control), Spa33_F67A/F70A_, Spa33_F70D_, MxiK_L46D_, or Spa33_F219D/Y221A_ with cartoon representation of sorting platform density observed in top panels. Positions of component proteins is shown corresponding to those in Figure 3d. f) Alternative model to Fig. 3, a-c, with two additional copies of Spa33 (purple) in place of the Spa33_short_ homodimers. The Spa33_N_ domains of the two additional Spa33 molecules extend into the lower pod density that is also occupied by MxiN (red).

## Discussion

Here we have used AlphaFold together with computational alanine scanning and *in vivo* mutant fitness characterization to create a model of the Spa33/MxiK portion of the *S. flexneri* sorting platform pod in which MxiK binds two copies of Spa33 via Spa33_N_, and each copy of Spa33 is complexed with a Spa33_short_ homodimer between Spa33_N_ and Spa33_C_ (Fig. 3a). This complex fits quite well into the upper portion of the experimental electron density map of the pod from cryo-electron tomography (cryo-ET). Notably, our proposed positioning of MxiK and Spa33 in the density map places these proteins closer to the inner membrane than is typically represented in sorting platform models (Fig. 3d, Suppl. Fig. 1)^5,10^. Here, the two Spa33_N_ domains occupy density that is generally shown as arising from MxiK, and MxiK shares the membrane-proximal density with MxiG^C^. However, this placement of the model gives better cross correlation values than any alternate positioning and visually fits the density map the best. Furthermore, our model explains the curious “donut hole” near the center of the pod electron density, as it arises from the space at the center of the Spa33/Spa33_short_ tetramer. The four Spa33_C_/Spa33_short_ domains fill the large section of density beneath the Spa33_N_ domains and the Spa33_C_ domains are positioned properly to bind the N-terminal peptide of MxiN from the region below. Finally, although we don’t have a good models of the MxiN homodimer structure, it is possible for MxiN to connect to Spa33_C_ at its position in our model of the upper pod (Fig. 3f).

The 2:1 Spa33:MxiK ratio in the pod is also congruent with the rapid exchange of Spa33 into and out of the sorting platform. FRAP experiments in *Yersinia* indicate that SctQ proteins are frequently cycling in and out of the sorting platform^7^, and this is likely the case in all T3SS. In the case of a 1:1 Spa33:MxiK ratio, the disengagement of a Spa33 molecule would temporarily eliminate a pod and potentially destabilize the sorting platform. The presence of a second copy of Spa33 also binding to MxiK would allow for one copy of Spa33 to exchange out without eliminating the pod connecting the spoke to the inner membrane. While this model of the Spa33/MxiK portion of the pod fits the density well and accounts for two copies of Spa33 in the pod, it leaves the lower portion of the pod density unmodeled. Part of this density must be due to the N-terminal regions of the two MxiN molecules, though it appears unlikely that MxiN alone can fill this volume (Fig. 3f).

The presence of significant density for the lower third of the pod without visible spokes in the Spa33/MxiK interaction mutant injectisomes is intriguing because this region of density would be expected to arise, at least in part, from the spoke protein MxiN. One possibility is that MxiN is present on these mutant pods but is bound in a more compact form (similar to Suppl. Fig. 7a) when the ATPase is absent. However, as shown in Fig. 3f, when the MxiN dimer is extended toward Spa47 it does not fill the density of the lower third of the pod. It is, therefore, likely that other protein molecules are present in this region.

The extra density in the lower portion of the pod could potentially be explained by an alternate model in which the two Spa33_short_ homodimers are replaced by the Spa33_C_ domains of two additional copies of Spa33. When asked to predict the multimer structure of two copies of Spa33, AlphaFold predicts a structure in which the Spa33_C_ domain of each Spa33 monomer is bound to the second copy in manner nearly identical to the Spa33_short_ homodimer (Suppl. Fig. 8), although the PAE values for this complex are much higher than those of the Spa33/Spa33_short_ structure. Replacement of the Spa33_short_ dimers in Fig. 3. a,b with additional copies of Spa33 gives a model for the pod that fits the upper density in the same orientation but now has an additional two Spa33_N_ domains available to fill the lower pod region (Fig. 4f). For this alternative model to be correct, the new Spa33_N_ domains must be reoriented relative to their position in Figure 4f to fit into the observed density and may make specific interactions with the MxiN dimer. A benefit of this alternate model is that it is consistent with previous studies that indicate there are four copies of SctQ present in each pod^7^. Due to the inaccuracy of AlphaFold MxiN structures, the need for conformational rearrangement of the additional Spa33_N_ domains, and their potential interactions with MxiN, we have not attempted to fit this alternative model into the lower pod density and leave that for future work.

## Materials and Methods

### AlphaFold predictions and computational alanine scanning

All predicted structures were made using the AlphaFold 3^23^ server (aphafoldserver.com) or ColabFold^24^. The top-ranked structure of each AlphaFold job was relaxed with Amber to correct residual steric clashes. Relaxed AlphaFold dimer structures were used as inputs for BUDE alanine scanning mutagenesis via the BAlaS server to identify mutationally sensitive interface residues (https://pragmaticproteindesign.bio.ed.ac.uk/balas/)^25^.

### Strains, plasmids, and mutagenesis

Δ*spa33* and Δ*mxiK S. flexneri* were obtained from C. Lesser (Massachusetts General Hospital, Cambridge, Massachusetts, USA) and A. Allaoui (Universite Libre de Bruxelles, Bruxelles, Belgium), respectively. Δ*spa33* and Δ*mxiK* strains were complemented by transformation with pUC18-derived pWPSF4 plasmid containing wild-type or mutant *spa33* and *mxiK* genes^18^. Mutants were created using overlap extension PCR and cloned into pWPSF4 using New England Biolabs HiFi Assembly Master Mix (NEB, E2621S). Defibrinated sheep red blood cells were purchased from Colorado Serum Company, Denver, Colorado.

### Contact-mediated hemolysis, CR-induced secretion, and western blotting

Δ*spa33* and Δ*mxiK S. flexneri* strains were transformed with pWPSF4 vectors carrying WT or mutant *spa33* or *mxiK* genes and grown overnight on selective tryptic soy agar plates. The next morning colonies were pickied, transferred to 10 mL tryptic soy broth (TSB). For contact-mediated hemolysis^18^, cultures were grown at 37° C to an OD_600_ of ∼0.6, centrifuged, and cell pellets resuspended in a volume of PBS to give an OD_600_ of 20. Three mL of defibrinated sheep red blood cells were diluted with 40 mL PBS, centrifuged, and resuspended in 3 mL of cold PBS. For each of five replicates, 50 μL of each *Shigella* sample were mixed with 50 μL of washed red blood cells in a single well in a round bottom 96 well plate, centrifuged for 10 minutes at 2000xg, and incubated for one hour at 37° C. 100 μL of PBs was then added to each sample and mixed thoroughly. Samples were then centrifuged again for 10 minutes at 2000xg. 100 μL of each sample supernatant was then removed and the absorbance at 545 nm was recorded. All A_545_ values were normalized to the average A_545_ of all WT samples.

For CR induction^19^, cultures were grown at 37° C to an OD_600_ ∼ 1.0, centrifuged, and cell pellets resuspended to an OD_600_ = 20 (approx.. 500 μL). CR was added to each cell suspension to a final concentration of 0.025% and cell suspensions incubated at 37° C for 20 minutes. Bacteria were then pelleted by centrifugation and supernatants aspirated to new tubes. Trichloroacetic acid was added to each supernatant sample to a concentration of 10% and samples incubated on ice for 30 minutes. Precipitated protein was pelleted by centrifugation at 14000xg for 15 minutes at 4° C. Precipitate pellets were washed twice with ice cold acetone. After removing acetone, pellets were dried by placing tubes in a 95° C heat block for 10 minutes. Dry pellets were resuspended in 40 μL PBS, 10 μL 5X SDS-PAGE loading buffer was then added, and samples boiled for 10 minutes. 20 μL of each sample was then separated on a 10% SDS-PAGE gel and blotted onto a nitrocellulose membrane. Each blot was treated with mouse serum containing anti-IpaB and anti-IpaD IgG antibodies, washed, and then incubated with anti-mouse secondary antibody labeled with Alexa Fluor 633. Bots were imaged on an iBright 1500 gel imaging station and band intensities quantified using ImageJ software.

To monitor protein expression with western blotting, a polyhistidine tag was added to the N-terminus of all MxiK variants, and to the N- and C-termini of all Spa33 variants in pWPSF4 and transformed into a Δ*mxiK* or Δ*spa33 S. flexneri* strain. T3SS expression was induced by growing all cells in 20 mL TSB at 37° C overnight with shaking at 200 rpm. The following morning, cell cultures were centrifuged for 10 minutes at 2000xg. Media supernatants were discarded, and cell pellets resuspended in 20 mL of histidine binding buffer (20 mM Tris, 500 mM NaCl, 15 mM imidazole, pH 8.0). Cells were lysed by brief sonication and lysate clarified by centrifugation at 30,000 xg for 30 minutes. Clarified lysate was passed over 1 mL of Ni-NTA resin, which was then washed with 20 column volumes of binding buffer. Captured his-tagged proteins were eluted in 3 mL of elution buffer (binding buffer with 500 mM imidazole). 25 μL of each eluted protein was mixed with 5 μL of 6X SDS-PAGE loading buffer and boiled for 5 minutes. 10 μL of each boiled sample was separated on a 12% SDS-PAGE gel. Gels were blotted onto a nitrocellulose membrane, blocked, and then incubated with mouse-derived anti-his IgG for 1 hour. Blots were then washed and incubated with an anti-mouse IgG labeled with Alexa Fluor 633 for 1 hour, washed again, and imaged on an iBright 1500 gel imager.

### Preparation of *Shigella* minicells for cryo-electron tomography (cryo-ET)

The sample preparation of *Shigella* minicells for cryo-ET approach was followed previous description^5,26,27^.*Shigella* mutants were streaked onto Congo red plates and incubated at 37℃ overnight. An isolated single colony was inoculated into 10 mL of tryptic soy broth (TSB) and grown overnight at 37 ℃ with shaking. To generate minicells, 2 mL of the overnight culture was inoculated into 200 mL of TSB and grown at 37 ℃ with shaking to late log phase. Kanamycin (50 μg/mL), ampicillin (100 μg/mL), and spectinomycin (100 μg/mL) were supplied into all bacterial plates and media for bacterial growth.

Minicells were purified by sequential centrifugation steps. The bacterial cultures were centrifuged twice at 2,000 x g for 10 min, followed by centrifugation of the supernatant at 20,000 x g for 10 min. Bacterial pellets were resuspended in 1mL of PBS, and then the resuspension was centrifuged at 2,000 x g for 5 min. The supernatant was then centrifuged at 15,000 x g for 5 min. The final bacterial pellet was resuspended in PBS.

Prior to vitrification, 10 nm gold tracer solution (Aurion) was added to the purified minicell sample. Then, 5 μL of the mixture was deposited on glow-discharged cryo-electron microscopy (cryo-EM) grids. To create a thin layer of cryo-fixation specimens, filter paper was placed on the cryo-EM grids, and specimens were then frozen in liquid ethane and propane mixture using a homemade gravity plunger and Leica EM GP2 plunger.

### Data collection using cryo-ET

The frozen specimens on cryo-EM grids were clipped by C-clip rings and C-clips (Thermo Fisher) and then loaded into a Titan Krios G2 microscope at Yale University. The microscope is equipped with a K3 summit direct detection camera and BioQuantum energy filter (Gatan). For imaging, the microscope was operated with 300kV acceleration voltage in almost -170 ℃ conditions. SerialEM was used to operate the microscope and acquire images^28^. For data collection by cryo-ET, the magnification for image recording was set at 42,000x, corresponding to 2.15Å physical pixel size, and the target defocus was set at ∼-4.9 μm. To collect tilt series images, the dose-symmetric scheme in FastTomo script was used to tilt the stage to cover an angle ranging from -48° to +48° in 3° increments^29^. The total electron dose for each tilt series was almost 65e^-^/Å^2^, and the dose fraction mode was used to collect 10 frames for each tilt angle image.

### Cryo-ET data processing and subtomogram averaging

Image drifting caused by electron beam exposure during image acquisition was corrected using MotionCor2^30^. IMOD was then used to generate image stacks and to trace 10 nm fiducial beads to align all images in each tilt series^30,31^.Gctf was used to estimate defocus values^32^, and then IMOD was used for contrast transfer function (CTF) correction. All tilt series were binned by a factor of 6 using the binvol function in IMOD^33^.

Tomo3D was used to reconstruct 6x6x6 binned tomograms using the simultaneous iterative reconstruction technique (SIRT)^34,35^. 693, 186, 167, and 97 tomograms were reconstructed from Spa33^F67A/F70A^ and Spa33^F70D^, MxiK^L46D^, and Spa33^F219D/Y221A^ mutants, respectively. From these tomograms, 1863, 527, 528, and 142 injectisomes were selected from Spa33^F67A/F70A^ and Spa33^F70D^, MxiK^L46D^, and Spa33^F219D/Y221A^ mutants, respectively.

For subtomogram averaging, 6x6x6 binned tomograms using weighted back projection (WBP) were reconstructed by Tomo3D, and i3 software was used to align the injectisome from each mutant^36,37^. For further structural analysis, unbinned subtomograms were extracted from tomograms based on the aligned position of the injectisome and subsequently binned by factors of 4 and 2 using the binvol function in IMOD. To compare structural phenotypes among the mutants, all injectisomes were aligned together using large masks. *In-situ* structures of the injectisome in each mutant were then determined based on the alignment positions of the injectisomes. Finally, each structure was refined using binned 2 subtomograms. For the fitting the predicted model, C6 symmetry was applied to a structure of the sorting platform structure of the wild type determined by binned 2 subtomograms to refine the structure. The sorting platform structure determined by unbinned subtomograms was then refined by focused refinement.

