## Supplemental Figures for "The pod components of the *Shigella* T3SS sorting platform accommodate multiple copies of Spa33 (SctQ)"

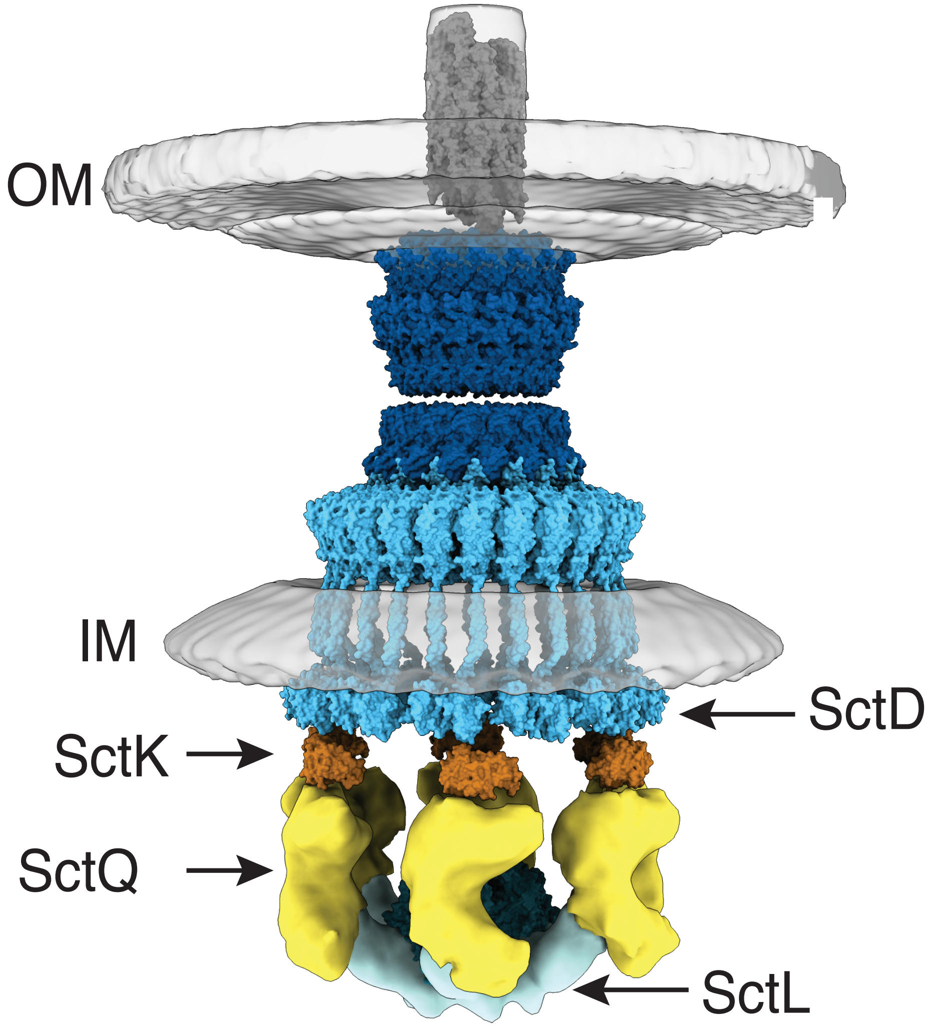


**Supplemental Figure 1:** Canonical model of the T3SS injectisome featuring extracellular needle extending from the outer membrane (OM), periplasm spanning basal body, and sorting platform on cytoplasmic side of inner membrane (IM). Positions of SP proteins are shown, including SctD cytoplasmic domain, SctK adaptor, SctQ pod protein, and SctL spoke.

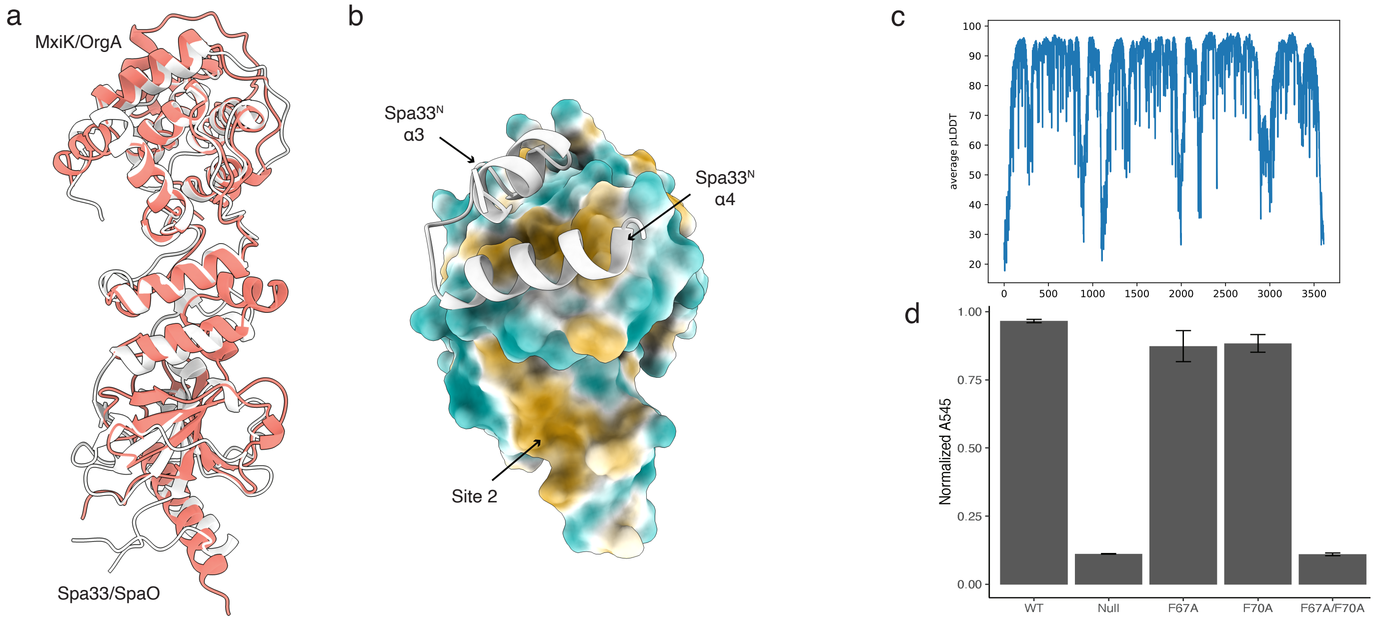


**Supplemental Figure 2:** a) Overlay of single Spa33/MxiK complex (white) with single SpaO/OrgA complex (red). b) Hydrophobic surface (gold is hydrophobic, cyan hydrophilic) of MxiK at the Spa33 interaction sites. The site1 interacting region of Spa33 is shown as white cartoon. Hydrophobic patch at binding site 2 is exposed here. c) pLDDT graph for the Spa33/Spa33/MxiK complex shown in main text Figure 1a. d) Contact mediated hemolysis data for Spa33 alanine mutants. Only the F67A/F70A double mutant has significant impact on hemoylysis.

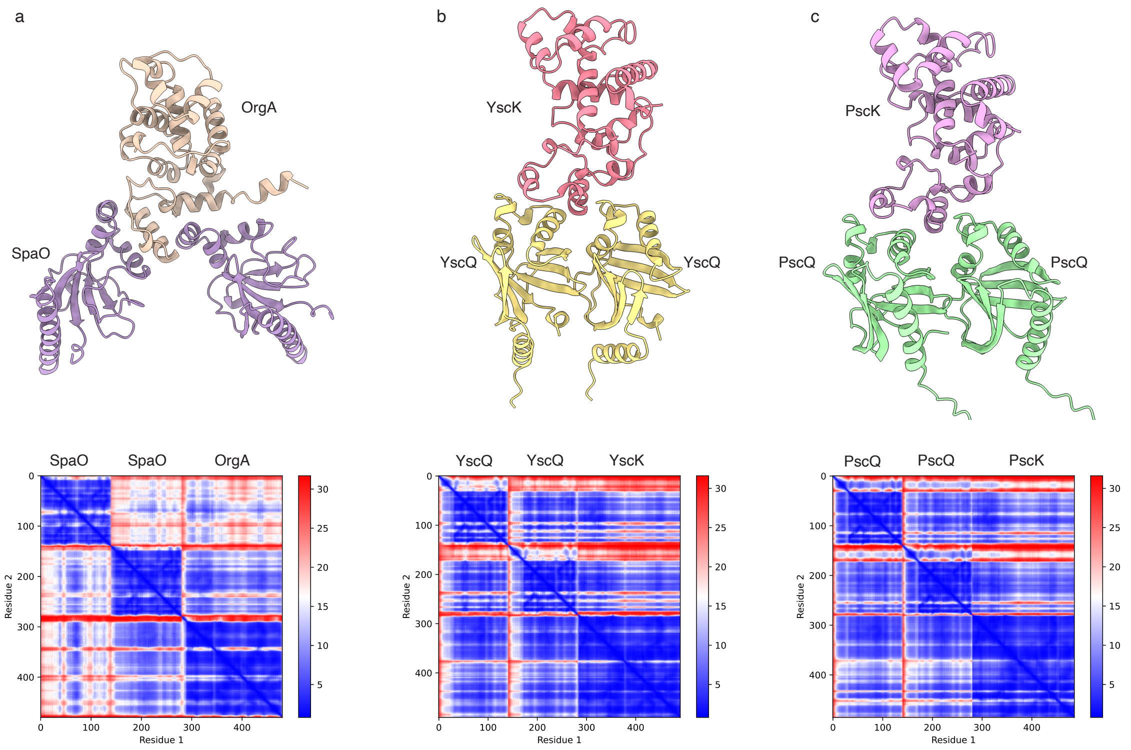


**Supplemental Figure 3:** AlphaFold generated 2SctQ:SctK complexes from a) *Salmonella*, b) *Yersinia*, and c)*Pseudomonas*, with corresponding PAE graphs below each cartoon.

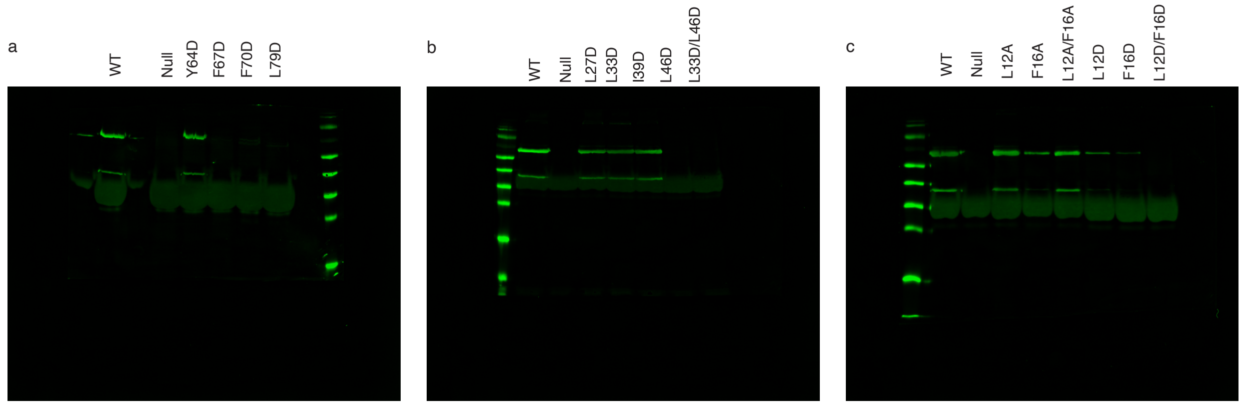


**Supplemental Figure 4:** Western blots of secreted IpaB and IpaD following 20 minute incubation of *Shigella flexneri* strains in PBS containing Congo Red. a) Spa33 mutants, b) MxiK site 1 mutants, and c) MxiK site 2 mutants. Quantification of these data are shown in main text Figure 1, g and h.

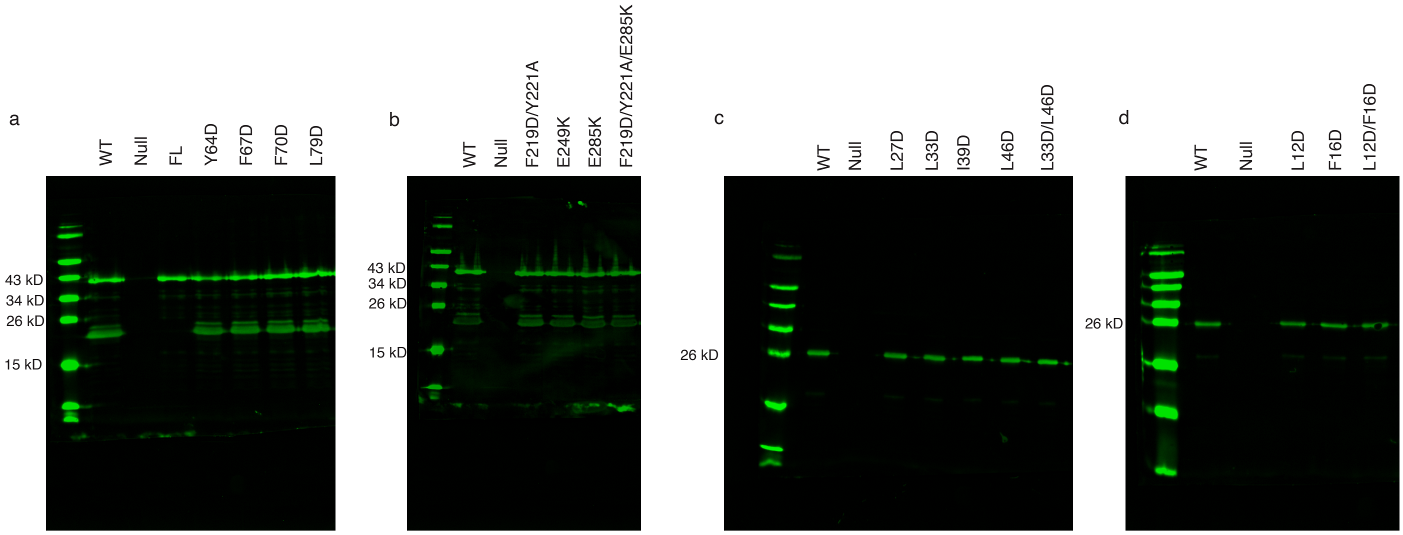


**Supplemental Figure 5:** Western blots using anti-His tage antibody of a) and b) His-Spa33-His variants; c) and d) His-MxiK variants. Null refers to uncomplemented Δ*spa33* or Δ*mxiK* cells. FL in panel (a) refers to a mutant of Spa33 in which the internal translation start site that produces $Spa33_{short}$ has been removed.

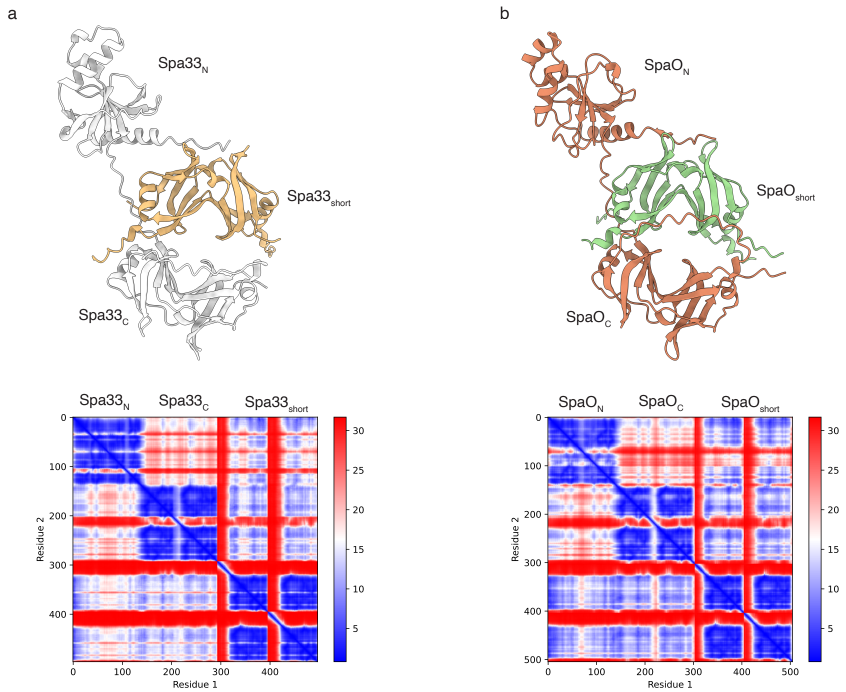


**Supplemental Figure 6:** Comparison of Spa33/Spa33$_{short}$ complex (a) with SpaO/SpaO$_{short}$ complex in (b), with corresponding PAE graphes below.

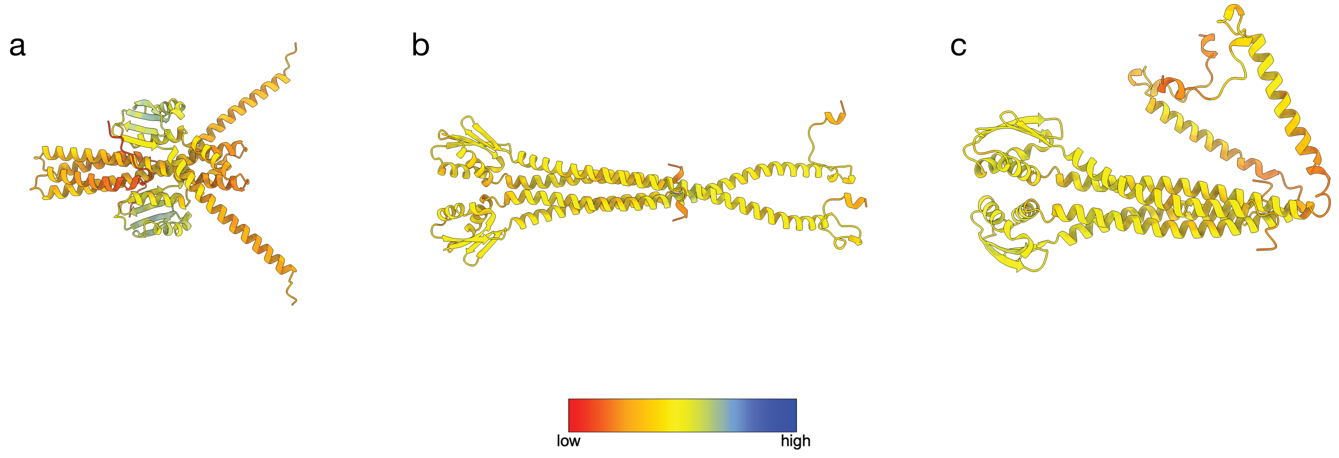


**Supplemental Figure 7:** Three separate AlphaFold predictions for the MxiN dimer structure, colored according to pLDDT scores, with red indicating very low confidence, yellow low confidence, and dark blue very high confidence.

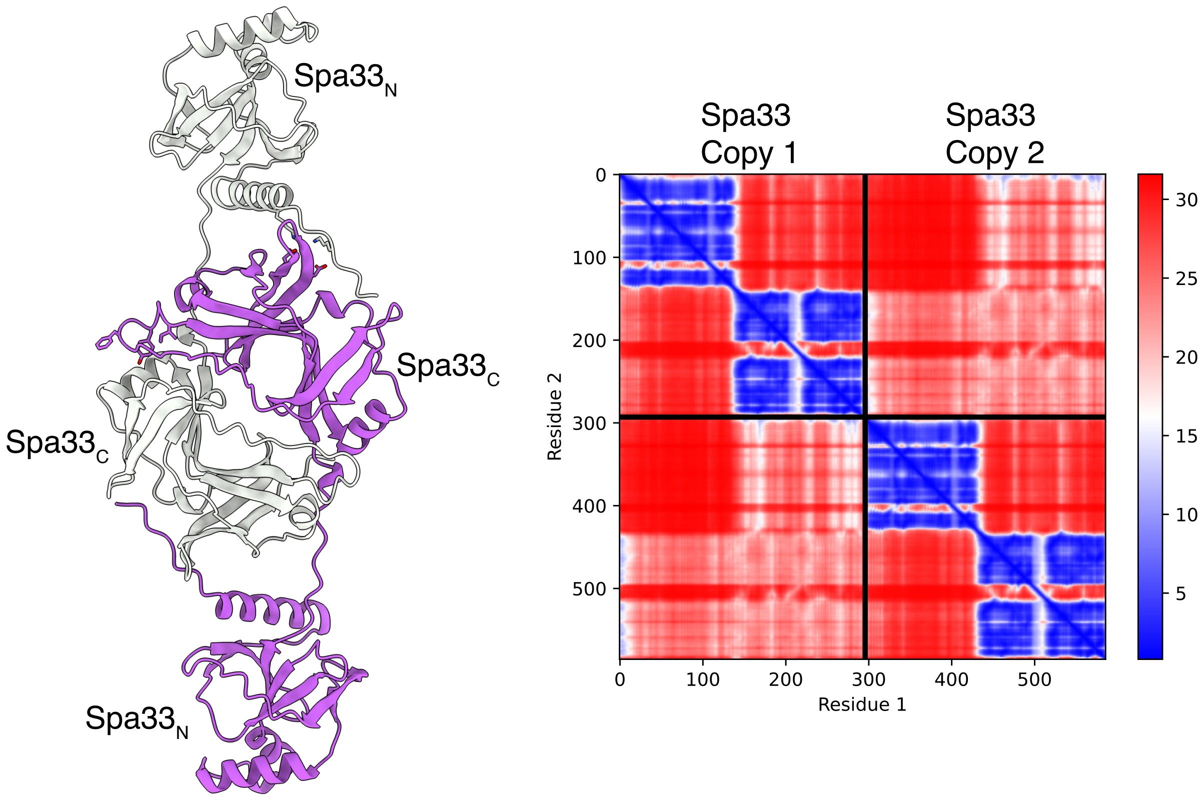


**Supplemental Figure 8:** AlphaFold structure for a Spa33 dimer with individual Spa33 molecules colored white and purple. In this model, the Spa33$_C$ domain of one Spa33 bind between the N- and C-terminal domains of the second Spa33 molecule in a manner nearly identical to the Spa33$_{short}$ dimer. PAE values (right) for the Spa33 dimer are much higher for intermolecular regions (off-diagonal blocks) than seen in the Spa33/Spa33$_{short}$ complex, indicating that AlphaFold is much less confident that its predicted Spa33 dimer is close to the “true” structure.
